# Methylomic signatures of tau and amyloid-beta in transgenic mouse models of Alzheimer’s disease neuropathology

**DOI:** 10.1101/2025.07.28.666515

**Authors:** Szi Kay Leung, Emma M. Walker, Stefania Policicchio, Aisha Dahir, Dorothea Seiler Vellame, Adam R. Smith, Rhian Swarbrick, Katie Lunnon, Emma L Dempster, Zeshan Ahmed, Eilis Hannon, Isabel Castanho, Jonathan Mill

## Abstract

Alzheimer’s disease (AD) is characterized by progressive neurodegeneration driven by tau and amyloid-β (Aβ) pathology. Emerging evidence implicates a role for epigenetic modifications, particularly altered DNA methylation (DNAm), in AD pathogenesis. However, few studies have comprehensively investigated DNAm in experimental models. Here, we profile DNAm dynamics in two widely used transgenic mouse models of tau (rTg4510) and Aβ (J20) neuropathology, focusing on variation in the entorhinal cortex and hippocampus. Using reduced representation bisulfite sequencing (RRBS) and methylation arrays across multiple disease stages, we identified widespread DNAm alterations associated with genotype and neuropathological burden. In rTg4510 mice, tau accumulation was linked to extensive DNAm remodeling at genes involved in neuronal plasticity and apoptosis (including *Dcaf5*, *Creb3l4*, and *As3mt*). J20 mice exhibited more modest changes annotated to immune-related genes, notably at *Grk2*, *Ncam2*, and *Prmt8*. Of note, tau-associated DNAm changes were more consistent across brain areas than those associated with Aβ pathology. Comparison with DNAm data from human studies revealed that a subset of DNAm sites mirrored those observed in the human AD cortex, including hypermethylation at *Ank1* and *Prdm16*. These findings provide evidence for pathology-associated epigenetic alterations in AD, highlight shared and distinct DNAm signatures of tau and Aβ, and offer insight into molecular mechanisms that may precede overt neurodegeneration. Our work underscores the utility of epigenomic profiling in transgenic models and provides a foundation for identifying novel targets for early intervention in AD.

## Introduction

Alzheimer’s disease (AD) – the most common type of dementia – is a chronic neurodegenerative disorder characterized by progressive memory deterioration, and cognitive and functional decline^1^. These symptoms result from progressive neuronal death synaptic loss, and brain atrophy^2^ associated with two core neuropathological processes: (1) the formation of neurofibrillary tangles (NFTs) as a result of intracellular aggregation of hyperphosphorylated tau protein, and (2) the development of amyloid plaques, which are extracellular deposits composed mainly of amyloid-β (Aβ) protein. While certain brain regions are relatively resistant to AD pathology, the hippocampus and entorhinal cortex – both involved in memory formation and recall – are particularly affected during the earliest stages of disease^3^. Despite significant progress in elucidating genetic risk factors for AD^4^, the precise molecular mechanisms that initiate and propagate NFT and Aβ pathology remain incompletely understood.

Emerging evidence suggests that epigenetic dysregulation plays a key role in the onset and progression of AD pathology. In particular, alterations in DNA methylation (DNAm) have been reported in postmortem human cortex in AD, implicating changes in the epigenome as a potential contributor to disease etiology and progression^5,6^. However, interpreting findings from such epigenome-wide association studies (EWAS) can be challenging due to confounding influences such as medication exposure and other secondary consequences of advanced disease, which complicate efforts to distinguish causal epigenetic changes from downstream effects of neurodegeneration. Transgenic (TG) rodent models of tau and Aβ pathology have helped identify mechanistic pathways involved in AD^7,8^. Recent transcriptomic studies in such models have revealed widespread gene expression changes driven by different AD-associated mutations^9^. To date, however, limited work has been undertaken to characterize epigenomic variation in AD-relevant mouse models^10^, with existing studies focusing on either global shifts in modifications to DNA or histones, or targeting a small number of candidate loci^11^. A comprehensive characterization of epigenetic alterations in AD-relevant mouse models could provide critical insights into early, mechanistic changes that precede overt late stages of disease and may reveal novel targets for therapeutic intervention.

In this study, we profiled DNAm in the entorhinal cortex, a region of the brain critically affected in early AD, using transgenic mouse models of tau and Aβ neuropathology. We profiled tissue at multiple time points from the rTg4510 transgenic model (ages 2, 4, 6, and 8 months) that expresses a human mutant form of the microtubule-associated protein tau (MAPT) gene^12^, and the J20 model (ages 6, 8, 10, and 12 months) that carries familial AD mutations in the amyloid precursor protein (APP) gene^13^. DNAm changes were quantified using two methodological approaches – reduced representation bisulfite sequencing (RRBS) and Illumina mammalian methylation array^14^ – with specific differences validated using bisulfite pyrosequencing. We subsequently compared the DNAm changes identified in the entorhinal cortex with DNAm profiles in the hippocampus, a region of the brain affected later in AD pathogenesis. Finally, we compared DNAm signatures identified in both models with data from a recent EWAS meta-analysis of postmortem human AD cortical tissue from >2,000 donors^15^. To our knowledge, this is the most comprehensive investigation of DNAm dynamics in AD-relevant transgenic mouse models to date, and the first to directly relate these changes to epigenetic alterations observed in human AD brains. Our findings provide further evidence for known responses to AD neuropathology and reveal novel genomic pathways associated with the accumulation of tau and Aβ.

## Results

### Profiling DNA methylation in mouse models of tau and Aβ neuropathology: experimental overview

We have previously described transcriptional changes in the entorhinal cortex and hippocampus associated with AD pathology in transgenic mice carrying human mutations in *MAPT (*rTg4510) and *APP* (J20)^16^. Building on our past work, we aimed to quantify site-specific DNAm changes associated with tau and Aβ neuropathology in the same animals (**Figure 1**). To achieve this, we used both highly-parallel reduced representation bisulfite sequencing (RRBS) and the Illumina mammalian methylation array to quantify genome-wide levels of DNAm in entorhinal cortex tissue dissected from rTg4510 TG and wild-type (WT) littermate controls (n = 31 WT, n = 30 TG, across ages 2, 4, 6, and 8 months) and J20 mice (n = 32 WT, n = 31 TG, ages 6, 8, 10, and 12 months) (**Figure 1**). Our RRBS experiments yielded a mean of 40.29M (SD = 6.14M) and 42.36M (SD = 7.24M) sequencing reads per sample in the rTg4510 and J20 models, respectively, with no significant differences in sequencing depth between TG and WT for either model (rTg4510: P = 0.908; J20: P = 0.449, **Table S1**). Following stringent preprocessing and quality control (see **Methods**), the final rTg410 entorhinal cortex DNAm dataset included a total of 1,364,030 sites (RRBS = 1,347,524 sites, array = 23,588 sites, overlap = 3,451 sites), while the J20 dataset included 1,388,735 sites (RRBS = 1,372,261 sites, array = 23,588 sites, overlap = 3,557 sites). As expected, these sites were primarily annotated to promoters given their high CpG content (rTg4510: n = 819,651 (60.8%), J20: n = 854,272 (62.3%), **Figure S1**). To assess consistency between platforms, we examined overlapping sites between RRBS and the array. Despite representing a small subset of total sites (15% of array sites, 0.26% of RRBS sites), DNAm estimates were highly correlated across technologies (rTg4510: r = 0.93; J20: r = 0.90; both P = 2.2E-16; **Figure S2**). Our analytical approach focused on two main analyses: (1) identifying group-level DNAm differences between TG and WT mice, and (2) testing associations between DNAm and quantitative measures of tau and Aβ pathology (assessed through immunohistochemistry in the same animals, **Figure 1**). We subsequently profiled DNAm in the hippocampal tissue dissected from the same animals using the Illumina mammalian methylation array to compare tau- and Aβ-associated differences across brain regions (**Figure 1**).

**Figure 1.**
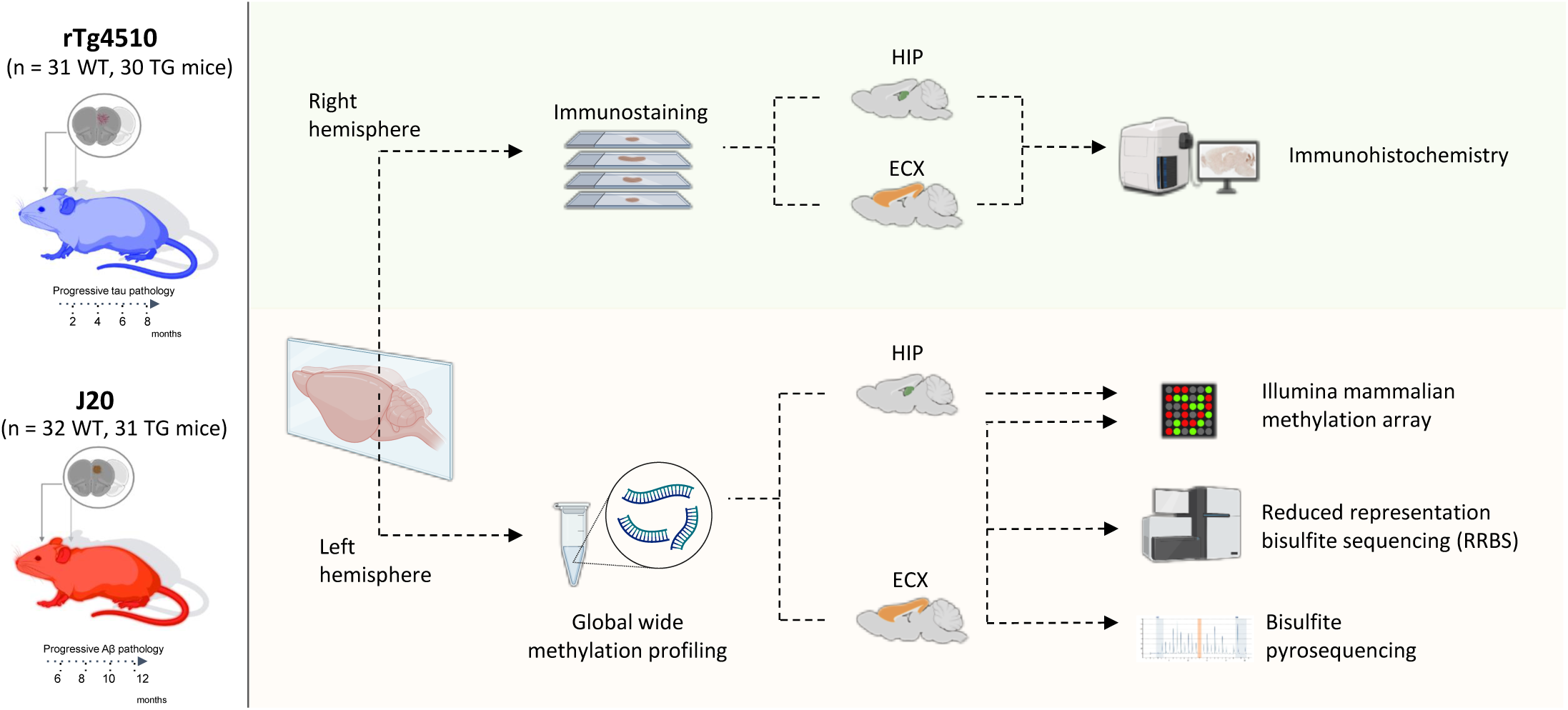
Study overview. To investigate DNA methylation signatures of tau pathology, we used the rTg4510 transgenic mouse line, which over-expresses human mutant (P301L) form of the microtubule-associated protein tau (*MAPT*). To investigate amyloid-β (Aβ) pathology, we used the J20 transgenic mouse line, which expresses a mutant (K670N/M671L and V717F) form of the human amyloid precursor protein (*APP*). Tissue was collected from transgenic (TG) and wild-type (WT) control mice at four time points. RRBS was used to quantify DNA methylation in the entorhinal cortex from both models. These data were complemented with DNA methylation data generated using the Illumina mammalian methylation array in both the entorhinal cortex and hippocampus from both models. Selected DMPs were validated using bisulfite pyrosequencing. ECX – entorhinal cortex, HIP – hippocampus, RRBS – Reduced representation bisulfite sequencing. Figure was created using BioRender.

### DNA methylation changes in rTg4510 cortex are annotated to genes regulating neuronal function and apoptosis

To identify differentially methylated positions (DMPs) in the entorhinal cortex associated with the *MAPT* transgene, we used a beta regression model to detect genotype-associated differences between rTg4510 TG and WT mice amongst sites profiled in the combined RRBS and array dataset (see **Methods**). In total, we identified 1,143 DMPs (false discovery rate (FDR) < 0.05) annotated to 514 genes between groups (**Figure 2A**, **Table S2**), with a significant enrichment of sites being hypermethylated in TG mice relative to WT mice (638 (55.8%) hypermethylated DMPs and 505 (44.2%) hypomethylated DMPs, binomial test: P = 9.29E-5). Several of the top-ranked DMPs between TG and WT mice were annotated to the *Mapt* gene itself (chr11:104318231, effect size = -1.04, FDR = 3.73E-30, **Figure S3**, **Table S2**) and a region covering *Prnp* that encodes prion protein readthrough transcript and prion protein (PrP) (n = 4 CpG sites, chr2:131910162 - chr2:131910201, mean effect size = 1.15, Empirical Brown’s method: P = 7.00E-5), which we validated using bisulfite pyrosequencing (**Figure S4**). These differences likely reflect the direct consequences of the transgene insertion^17^ and mirror differential expression of the same genes identified in our previous study^9^. The list of DMPs also included sites annotated to other genes known to be affected by the CaMKIIα-tTA transgene insertion (including DMPs annotated to *Ncapg2* and *Ptprn2*) and *MAPT* (including DMPs annotated to *Fgf14*) transgenes^17^ (**Figure S3**).

**Figure 2.**
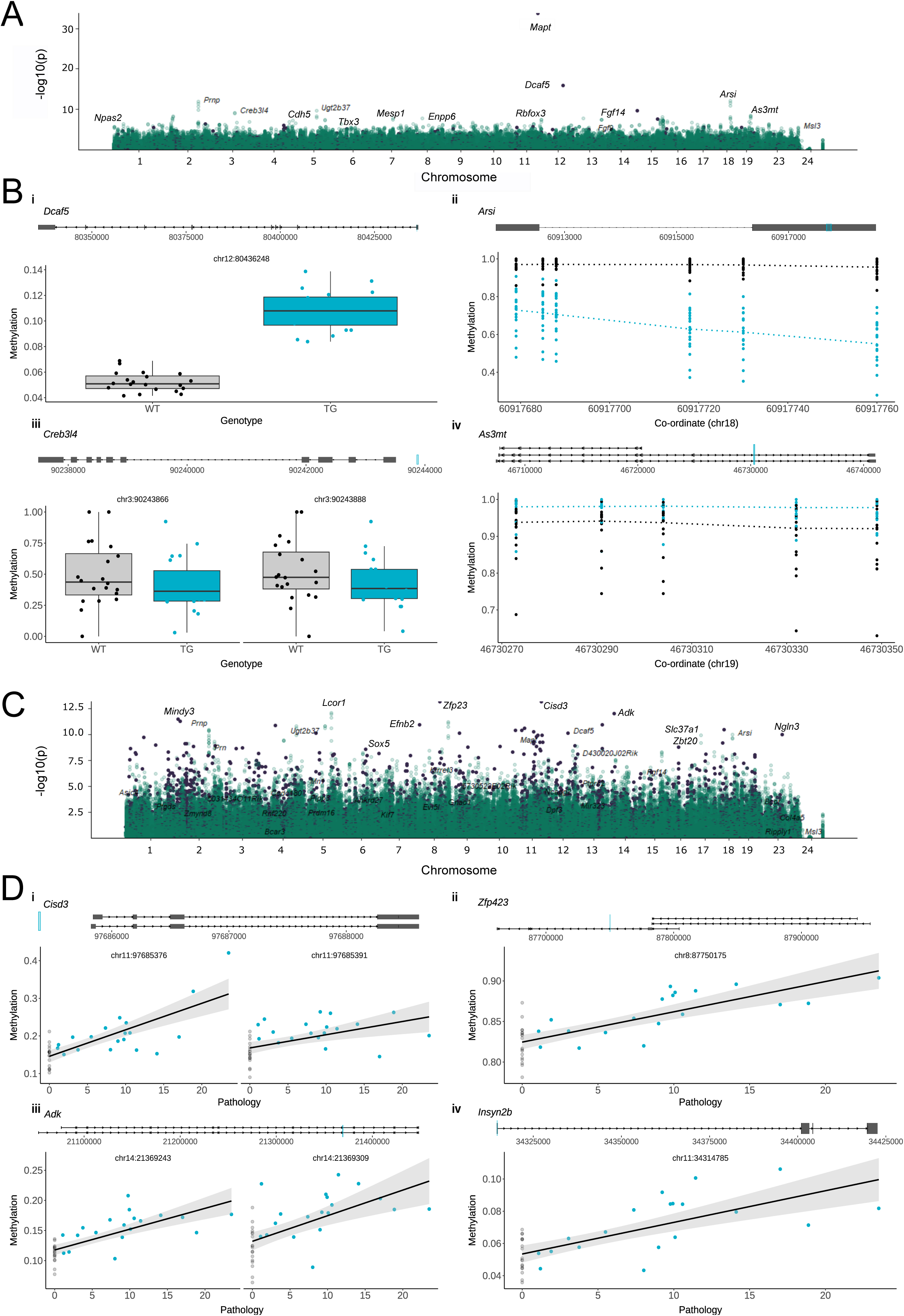
Differential DNA methylation in rTg4510 mice. **(A)** Manhattan plot showing transgene-associated differential methylation positions (DMPs) in rTg4510 mice (n = 61, 31 WT, 30 TG). Purple and green dots refer to sites identified using the Illumina mammalian DNA methylation array and RRBS, respectively. **(B)** Shown are the gene tracks and top-ranked differentially methylated sites annotated to **(i)** *Dcaf5* (chr12:80436248, effect size = 0.39, FDR = 1.3E-12), **(ii)** *Arsi* (n = 6 CpG sites, chr18:60917760 - chr18:60917679, mean effect size = -1.51), **(iii)** *Creb3l4* (n = 2 CpG sites, chr3:90243866 - chr3:90243888, mean effect size = -2.34), and **(iv)** *As3mt* (n = 5 CpG sites, chr19:46730273 - chr19:46730349, mean effect size = 1.48), between WT and rTg4510 TG mice. Black and blue dots refer to WT and TG respectively, and the blue lines on the tracks refer to the location of DMPs relative to the gene. **(C)** Manhattan plot showing DMPs associated with levels of tau measured using immunohistochemistry in rTg4510 TG mice. Purple and green dots refer to sites identified using the Illumina mammalian DNA methylation array and RRBS, respectively. **(D)** Shown are the gene tracks and top-ranked differentially methylated sites annotated to **(i)** *Cisd3* (chr11:97685376, effect size = 0.025, FDR = 7.84E-10; chr11:97685391, effect size = 0.013, FDR = 4.32E-3), **(ii)** *Zfp23* (chr8:87750175, effect size = 0.017, FDR = 7.84E-10) **(iii)** *Adk* (chr14:21369243, effect size = 0.016, FDR = 7.19E-9; chr14:21369309, effect size = 0.018, FDR = 2.87E-4), **(iv)** *Insyn2b* (chr11:34314785, effect size = 0.015, FDR = 6.65E-8), associated with progressive tau pathology in rTg4510 TG mice. Black and blue dots refer to WT and TG respectively, and the blues lines on the tracks refer to the location of CpG sites.

Beyond differences at these sites, we found highly significant genotype-associated differences in DNAm at sites and groups of sites annotated to many genes, including *Dcaf5* (chr12:80436248, effect size = 0.39, FDR = 1.3E-12), *Arsi* (n = 6 CpG sites, chr18:60917760 - chr18:60917679, mean effect size = -1.51, Empirical Brown’s method: P = 4.64E-6), *Creb3l4* (n = 2 CpG sites, chr3:90243866 - chr3:90243888, mean effect size =-2.34, Empirical Brown’s method: P = 5.03E-05), and *As3mt* (n = 5 CpG sites, chr19:46730273 - chr19:46730349, mean effect size = 1.48, Empirical Brown’s method: P = 3.46E-4) (**Figure 2B**, **Table S2**). *Dcaf5* is a substrate receptor of the CUL4-DDB1 E3 ubiquitin-protein ligase (CRL4) complex, which binds to the cytosolic region of APP and is involved in the regulation of the ubiquitin-proteasome system^18^. *Arsi* encodes arylsulfatase family member I (ARSI), a paralogue of arylsulfatase B (ARSB), which has been associated with AD^19^ and Parkinson’s disease^20^, and reported in an EWAS of dementia with Lewy bodies (DLB)^21^. *Creb3l4* encodes a CREB (cAMP responsive element binding) transmembrane protein that functions as transcriptional activator of neuronal plasticity and protection. Of relevance, studies have shown that CREB signaling is down-regulated in AD brains^22^, CREB levels have been reported to be decreased in hippocampal neurons from Tg2576 mice^23^, and CREB has been suggested to negatively regulate the transcription of tau^24^. *As3mt*, encoding for arsenite methyltransferase, has previously been implicated in schizophrenia with a role in neuronal development and differentiation, and was also recently identified to be hypermethylated in AD donors with psychosis^25^.

Functional enrichment analysis of the genes annotated to genotype-associated DMPs (see **Methods**) revealed a strong enrichment for pathways that are highly relevant to tau pathology including the positive regulation of apoptosis (FDR = 5.76E-20) and the activity of enzymes involved in the metabolism of inositol (e.g. inositol tetrakisphosphate 1-kinase activity (FDR = 1.11E-21)) and sphingosine (e.g. sphingosine N-acyltransferase activity (FDR = 8.19E-17)) (**Table S3**), which have been previously implicated in AD and are involved in mediating cell death^26,27^. Of note, among human phenotype ontologies enriched amongst genes annotated to DMPs were neurofibrillary tangles (FDR = 4.3E-9) and global brain atrophy (FDR = 1.74E-09) (**Table S3**). Taken together, we detected significant changes in epigenetic regulation of key genes that play an important role in neuronal survival and plasticity associated with tau neuropathology, with P301L tau overproduction inducing neuronal cell death.

We next sought to explore DNAm changes associated with the progression of neuropathology in TG mice by identifying sites at which DNAm levels were associated with levels of tau in the entorhinal cortex measured using immunohistochemistry (see **Methods**). In total, we identified 7,499 DMPs (FDR < 0.05) annotated to 2,789 genes (**Figure 2C**, **Table S2**) associated with increased levels of tau, with the top-ranked DMP being located upstream of the *Cisd3* gene (chr11:97685376, FDR = 7.84E-10, **Figure 2D**) encoding an iron–sulfur domain-containing protein that plays a vital role in controlling mammalian lifespan and has been shown to mediate AD-associated neuronal loss^28^. Other top-ranked sites were located in an intron of *Zfp423* (chr8:87750175, effect size = 0.017, FDR = 7.84E-10), and of *Adk* (chr14:21369243, effect size = 0.016, FDR = 7.19E-9) and in the promoter regions of *Insyn2b* (chr11:34314785, effect size = 0.015, FDR = 6.65E-8) (**Figure 2D**). Of note, *Zfp423* encodes a zinc finger transcription factor that regulates apolipoprotein D^29^ and is upregulated in neurons from the dentate gyrus in aged and AD mice^30^. To refine these sites associated with tau pathology, we used a beta regression model to further identify sites at which DNAm was characterized by a significant interaction between genotype and age (**Table S2**). A total of 541 DMPs, mapping to 372 genes, were significant in both analyses (FDR < 0.05), highlighting loci where variable DNAm is strongly associated with progression of tau pathology in TG mice (**Table S2**).

### DNA methylation changes in J20 cortex are annotated to genes involved in mitochondrial homeostasis

To investigate DNAm changes associated with mutations in *APP*, we applied the same analysis to genome-wide DNAm data from the entorhinal cortex of J20 mice. We identified 2,521 genotype-associated DMPs (FDR < 0.05) annotated to 920 genes (**Figure 3A**, **Table S4**).

**Figure 3.**
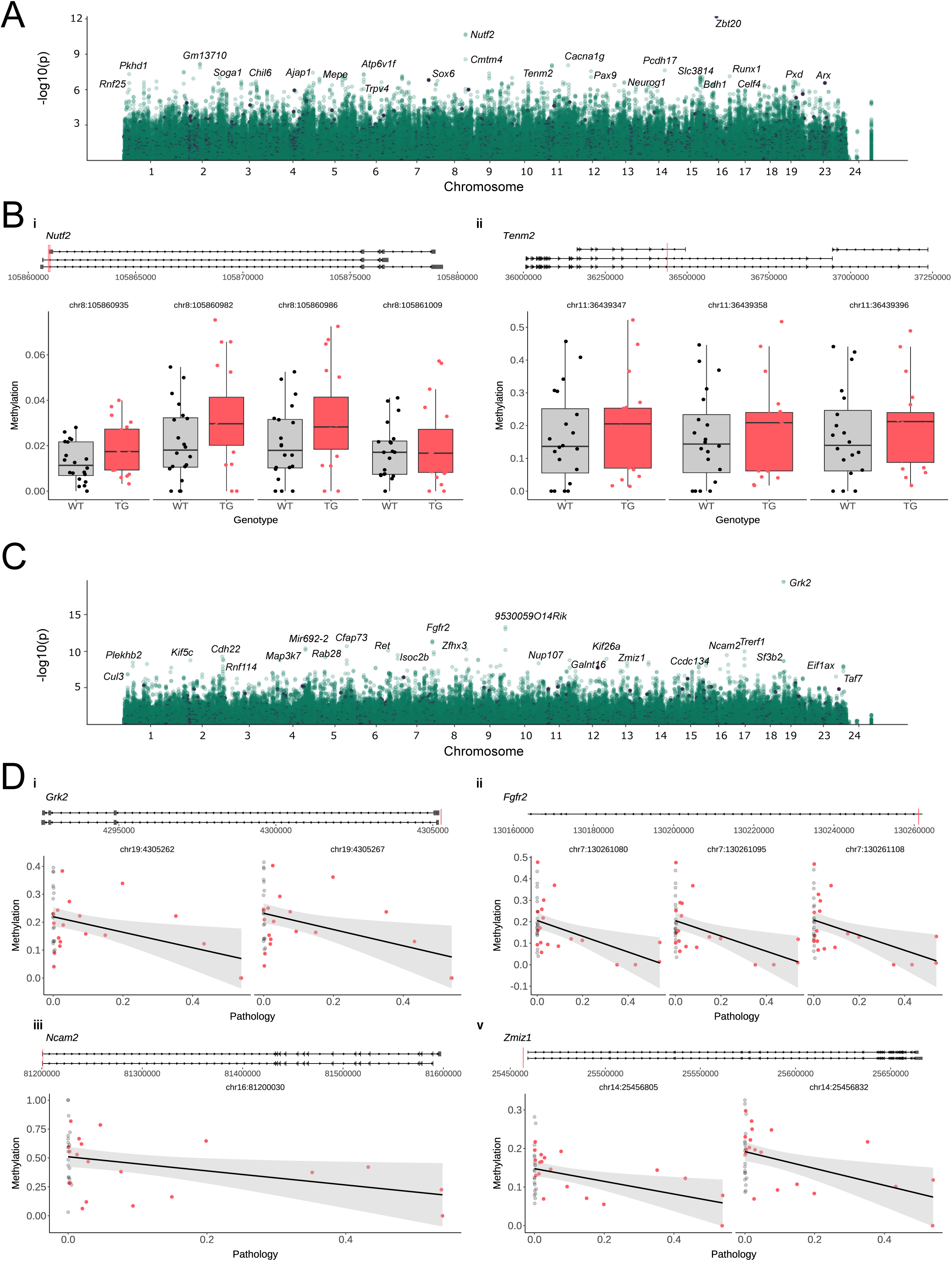
Differential DNA methylation in J20 mice. **(A)** Manhattan plot showing transgene-associated differential methylation positions (DMPs) in J20 mice (n = 63, 32 WT, 31 TG). **(B)** Shown are the gene tracks and top-ranked differentially methylated sites annotated to **(i)** *Nutf2* (n = 4 CpG sites, chr8:105860982 - chr8:105861009, mean effect size = 1.73. Top-ranked site: chr8:105860982, effect size = 2.19, FDR = 1.72E-5) and **(ii)** *Tenm2* (n = 3 CpG sites, chr11:36439347 - chr11:36439396, mean effect size = 2.93. Top-ranked site: chr11:36439396, effect size = 2.92, FDR = 1.54E-3) between WT and J20 TG mice. Black and red dots refer to WT and TG respectively, and the red lines on the tracks refer to the location of CpG sites. **(C)** Manhattan plot showing differential methylation positions (DMPs) associated with progressive Aβ pathology in J20 TG mice. **(D)** Shown are the gene tracks and top-ranked differentially methylated sites annotated to **(i)** *Grk2* (chr19:4305267, effect size = -2.02, FDR = 2.61E-14; chr19:4305262, effect size = -1.99, FDR = 2.61E-14), **(ii)** *Fgfr2* (chr7:130261095, effect size = -1.661, FDR = 1.10E-6; chr7:130261108, effect size = -1.657, FDR = 1.23E-6), **(iii)** *Ncam2* (chr16:81200030, effect size = -2.01, FDR = 2.67E-5), and **(iv)** *Zmiz1 (*n = 2 DMP sites, chr14:25456805-chr14:25456832, mean effect size = -1.31, P = 8.07E-6), between WT and J20 TG mice with progressive Aβ pathology. Black and red dots refer to WT and TG respectively, and the red lines on the tracks refer to the location of CpG sites.

Similar to our findings in rTg4510 mice, there was a significant enrichment of hypermethylated sites in TG mice (1,785 (70.8%) hypermethylated DMPs and 736 (29.2%) hypomethylated DMPs, binomial test: P = 1.11E-99). The top-ranked J20 DMP was annotated to an intronic region of *Zbtb20*, a gene known to be disrupted by the transgene insertion^13^, suggesting this methylation change likely reflects a direct effect of the genetic perturbation. Other notable DMPs (**Table S4**) included sites annotated to *Nutf2* (n = 4 CpG sites, chr8:105860982 - chr8:105861009, mean effect size = 1.73, Empirical Brown’s method: P = 9.99E-6, **Figure 3B**) – encoding nuclear transport factor 2 (NUTF2), which is reported to aggregate in hippocampal neurons in AD^31^ – and *Tenm2* (n = 3 CpG sites, chr11:36439347 - chr11:36439396, mean effect size = 2.93, Empirical Brown’s method: P = 2.57E-4, **Figure 3B**) – encoding teneurin transmembrane protein 2, which is involved in neural development and associated with AD pathogenesis^32^. Functional enrichment analysis of the genes annotated to DMPs associated with J20 genotype using GREAT (see **Methods**) identified numerous pathways and functions related to AD including many related to neuroinflammation (e.g. chemokine activity (FDR = 5.81E-10), the cytolytic granule (FDR = 1.37E-15) and cell chemotaxis (FDR = 3.54E-13) (**Table S3**).

We next examined DNAm changes associated with Aβ levels quantified in the entorhinal cortex using immunohistochemistry (see **Methods**) identifying 2,136 DMPs (FDR < 0.05) annotated to 754 genes (**Figure 3C**, **Table S4**). The two top-ranked DMPs (chr19:4305267, effect size = -2.02, FDR = 2.61E-14; chr19:4305262, effect size = -1.99, FDR = 2.61E-14, **Figure 3D**) were annotated to the first intron of *Grk2*, which encodes a G protein-coupled receptor kinase that has been widely implicated in the generation of Aβ in AD^33^, with its overexpression recognized as a primary hallmark of mitochondrial lesions^34^. Other top-ranked sites were annotated to the promoter of *Fgfr2* (chr7:130261095, effect size = -1.661, FDR = 1.10E-6; chr7:130261108, effect size = -1.657, FDR = 1.23E-6, **Figure 3D**). A number of these DMPs (n = 38) were also characterized by a significant genotype-by-age interaction effect (**Table S4**), and included sites annotated to the promoter region of *Ncam2* (chr16:81200030, effect size = -2.01, FDR = 2.67E-5) encoding neural cell adhesion molecule 2 (NCAM2), an intergenic region of *Zmiz1* (n = 2 DMP sites, chr14:25456805-chr14:25456832, mean effect size = -1.3, P = 8.07E-6), and the promoter regions of *Prmt8* (n = 2 DMP sites, chr6:127768697 - chr6:127768710, mean effect size = -0.95, Empirical Brown’s method: P = 9.56E-4) encoding protein arginine methyltransferase 8 (PRMT8) (**Figure 3D**). Of note, levels of NCAM2 have been found to be depleted in the hippocampus of AD donors and APP transgenic mouse models^35^. Recent studies identified ZMIZ1 as significantly associated with mitophagy in Lewy body disease and reduced neuropathology burden, indicating a neuroprotective role for facilitating appropriate degradation of damaged mitochondria^36^. PRMT8 is reported to modulate mitochondrial bioenergetics and neuroinflammation in response to hypoxic stress^37^. Collectively, our results suggest that DNA methylation changes annotated to genes involved in neuroinflammation and the regulation of mitochondrial function are associated with progressive Aβ pathology.

Finally, we compared DNAm differences identified in J20 mice with those identified in rTg4510 mice at sites profiled in both models. Overall the correlation of effect sizes between models was low (genotype: RRBS sites r = 0.03, array sites r = 0.04; pathology: RRBS sites = 0.04, array sites = 0.18). These results suggest that there is limited overlap in DNAm changes associated with tau and Aβ pathology, corroborating results from our previous gene expression study^16^. Six genotype-associated DMPs (**Table S5**), however, were identified in both transgenic models, including four sites annotated to the promoter of the *SATB1* gene associated with genotype (TG vs WT) differences. Notably, SATB1 is a DNA binding protein associated with neurodegenerative disease that functions to prevent cellular senescence in post-mitotic dopaminergic neurons^38,39^.

### Tau and Aβ-associated DNA methylation changes are more pronounced in the hippocampus

We next performed DNAm profiling on hippocampal tissue from the same rTg4510 and J20 mice using the Illumina mammalian methylation array (rTg4510 WT = 31, TG = 30; J20 WT = 32, TG = 31; see **Methods** for details). Following preprocessing and stringent quality control, we retained all 23,588 array sites in the hippocampus dataset that overlapped with those profiled in the entorhinal cortex. In the rTg4510 model, we observed more extensive DNAm differences in the hippocampus than in the entorhinal cortex, of which 374 DMPs were associated with genotype differences between TG and WT (**Figure 4A**, **Table S6**), and 8,857 DMPs were associated with immunohistochemically measured tau pathology (**Figure 4A**, **Table S6**).

**Figure 4.**
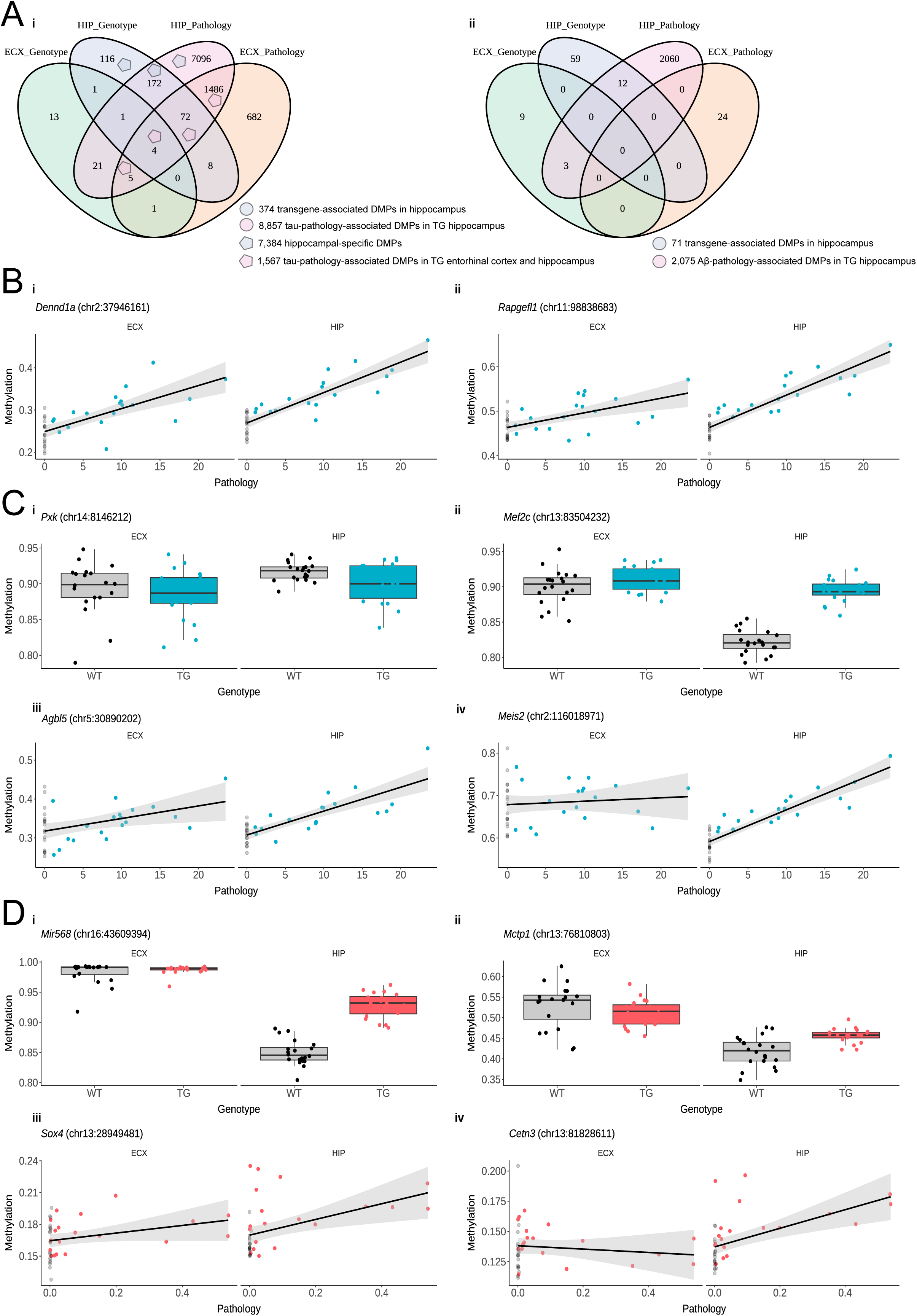
Transgene-associated DMPs exhibit distinct DNA methylation profiles in the entorhinal cortex and hippocampus. **(A)** Venn diagram showing the number of array-detected significant DMPs detected in **(i)** rTg4510 and **(ii)** J20 entorhinal cortex and hippocampus associated with genotype and progressive pathology. **(B)** Shown are the top-ranked DMPs associated with progressive tau pathology in rTg4510 TG mice in both the entorhinal cortex and the hippocampus: **(i)** *Dennd1a* and **(ii)** *Rapgefl1*. Black and blue dots refer to WT and TG respectively. **(C)** Shown are the top-ranked DMPs associated with tau pathology in rTg4510 TG vs WT mice in the hippocampus but not in entorhinal cortex: **(i)** *Pxk* (chr14:8146212; hippocampus: effect size = -0.39, FDR = 2.73E-06; entorhinal cortex: effect size = -0.14, FDR = 0.89), **(ii)** *Mef2c* (chr13:83504232 - hippocampus: effect size = 0.34, FDR = 7.14E-06; entorhinal cortex: effect size = -0.18, FDR = 0.506). Shown are also the top-ranked DMPs associated with associated with progressive tau pathology in rTg4510 mice in the hippocampus but not in entorhinal cortex: **(iii)** *Agbl5* (chr4:109806599 - hippocampus: effect size = 0.012, FDR = 1.05E-40; entorhinal cortex: effect size = 0.0086, FDR = 0.052), **(iv)** *Meis2* (chr2:116018971 - hippocampus: effect size = 0.014, FDR = 4.44E-25; entorhinal cortex: effect size = 0.0014, FDR = 0.84). Black and blue dots refer to WT and TG respectively. **(D)** Shown are the top-ranked DMPs associated with Aβ pathology in J20 TG vs WT mice in the hippocampus but not in entorhinal cortex: **(i)** *Mir568* (chr16:43609394; hippocampus: effect size = 0.80, FDR = 1.46E-05; entorhinal cortex: effect size = -0.23, FDR = 0.78) and **(ii)** *Mctp1* (chr13:76810803; hippocampus: effect size = 0.33, FDR = 6.63E-03; entorhinal cortex: effect size = -0.12, FDR = 0.74) associated with Aβ pathology in J20 TG vs WT mice. Shown are also the top-ranked DMPs associated with progressive Aβ pathology in J20 mice, significant in the hippocampus but not in the entorhinal cortex: **(iii)** *Sox4* (chr13:28949481; hippocampus: effect size = 0.013, FDR = 1.79E-11; entorhinal cortex: effect size = 0.15, FDR = 0.96), **(iv)** *Cetn3* (chr13:81828611; hippocampus: effect size = 0.013, FDR = 1.79E-11; entorhinal cortex: effect size = 0.15, FDR = 0.96). Black and red dots refer to WT and TG respectively.

Although genotype-associated differences showed minimal concordance between hippocampus and entorhinal cortex across all sites tested (r = 0.03, P = 1.09E-6, **Figure S5**), six DMPs (*Dcaf5* (chr12:80436248), *Satb1* (chr17:51746925), *Cltc* (chr11:8670046), *Mapt* (chr11:104318231), *Ncapg2* (chr12:116425797), *Fgf14* (chr14:124676565)) were shared between regions (**Figure S6**) and their corresponding effect sizes were completely correlated (r = 1, P = 3.37E-5, **Figure S5**). In contrast, tau pathology–associated DMPs were more strongly correlated across brain regions (r = 0.42, P = 0, **Figure S5**), with 1,567 DMPs shared between the entorhinal cortex and hippocampus (**Figure 4A, 4B**). Among the 7,384 significant hippocampal-specific DMPs (genotype: n = 288 sites annotated to 238 genes, tau pathology: n = 7,268 sites annotated to 2,487 genes), the top-ranked DMPs included sites annotated to the exonic region of *Pxk* (chr14:8146212; hippocampus: effect size = -0.39, FDR = 2.73E-06; entorhinal cortex: effect size = -0.14, FDR = 0.89), *Mef2c* (chr13:83504232 - hippocampus: effect size = 0.34, FDR = 7.14E-06; entorhinal cortex: effect size = -0.18, FDR = 0.51), and *Meis2* (chr2:116018971 - hippocampus: effect size = 0.014, FDR = 4.44E-25; entorhinal cortex: effect size = 0.0014, FDR = 0.84) (**Figure 4C, Table S6**). Of note, MEIS2 downregulation has been found to improve memory retention of AD mice^40^ and MEF2C is a synaptic transcription factor implicated in AD^41^. In the J20 model, we also observed more DMPs in the hippocampus than in the entorhinal cortex (**Figure 4A**), with 71 genotype-associated DMPs and 2,075 DMPs associated with levels of Aβ (**Figure 4D, Table S7**). In contrast to the rTg4510 model, however, no DMPs were shared between brain regions and overall effect sizes for Aβ-associated methylation differences across all profiled sites were uncorrelated across tissues (**Figure S7**). These results suggest that tau-associated DNAm differences in rTg4510 mice are more similar across brain regions, with more widespread methylation changes in the hippocampus than the entorhinal cortex, compared to Aβ-related changes in J20 mice that appear to be highly region-specific.

### Accumulation of tau is associated with accelerated epigenetic age

Given that recent studies have reported an association between accelerated DNAm age and neuropathology in human AD^42^, we next applied epigenetic clock algorithms to our data. Using an epigenetic clock derived from sites on the Illumina methylation array previously calibrated on mouse cortex^43^, we found an overall strong relationship between epigenetic age and chronological age across all samples (TG and WT) in both the entorhinal cortex (rTg4510: r = 0.89, P = 1.52E-14; J20: r = 0.81, P = 2.1E-10) and hippocampus (rTg4510: r = 0.87, P = 7.39E-14; J20: r = 0.90, P = 4.65E-15) (**Figure S8**). We then investigated whether there was an acceleration of epigenetic age associated with AD pathology, and observed an increase in epigenetic age specifically in the hippocampus of older TG rTg4510 compared to WT mice (6 months: t-test P = 0.031; 8 months: t-test P = 0.032); no epigenetic age acceleration was observed in the entorhinal cortex, potentially reflecting the more extensive DNAm changes observed in the hippocampus relative to the cortex. TG J20 mice did not show an increase in epigenetic age compared to WT mice (**Figure S8)**, supporting previous studies on human postmortem tissue that found accelerated epigenetic age was more positively associated with neurofibrillary tangles than amyloid load^44^.

### Changes in DNA methylation parallel those identified in human Alzheimer’s disease brains

To evaluate the translational relevance of DNA methylation (DNAm) changes observed in transgenic models of AD, we compared our findings with those from a recent large-scale epigenome-wide association study (EWAS) of postmortem human cortex (n > 2,000 donors)^15^, which identified 334 differentially methylated positions (DMPs) associated with Braak neurofibrillary tangle (NFT) stage. Of the 171 genes annotated to these human DMPs, 149 had conserved mouse orthologs, and notably of these, 38 (25.5%) were also annotated to sites significantly associated with either genotype or tau pathology progression in the rTg4510 mouse model (**Figure 5A**, **Table S8**). The most significant DMPs in the rTg4510 entorhinal cortex overlapping with those identified in AD brain were located in the intronic region of *Ank1* (**Figure 5B**) spanning three sites that were significantly hypermethylated in TG mice (chr8:23023240: effect size = 0.905, FDR = 1.03E-02; chr8:23023210: effect size = 0.912, FDR = 1.37E-02, chr8:23023192: effect size = 0.926, FDR = 1.73E-02). We independently validated this finding by quantifying methylation of a subset of the *Ank1*-annotated sites using bisulfite pyrosequencing (chr8:23023192: t-test = 8.28E-3; chr8:23023240: t-test = 1.3E-2) (**Figure 5B**). Of note, hypermethylation of sites annotated to the *ANK1* gene represents one of the most robust findings in studies of human AD cortex^45^. Similarly, in the J20 mouse model of amyloid pathology, 20 genes (13.4%) annotated to human AD-associated DMPs also exhibited significant methylation changes associated with either genotype or Aβ pathology progression. The most significant overlap was observed at a CpG site in the promoter of *Tspan14* (chr14:40966816, effect size = -1.49, FDR = 1.16E-02).

**Figure 5.**
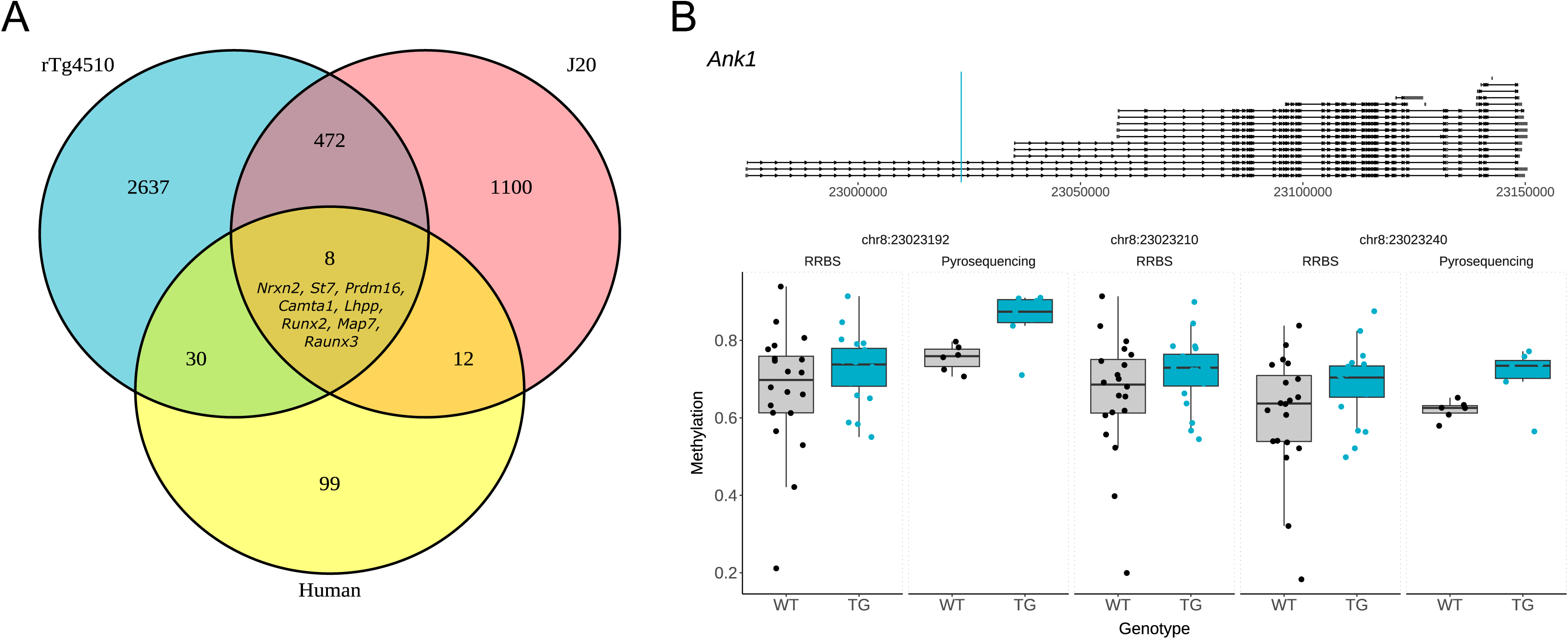
DNA methylation differences identified in mice overlap with AD-associated methylation changes in human brains. **(A)** Venn diagram showing the number of genes annotated to significant DMPs identified in rTg4510 mice, J20 mice and human AD. DMPs associated with 8 genes overlapped between both models and human AD. **(B)** Scatter plot showing the methylation estimates from RRBS (chr8:23023240: effect size = 0.905, FDR = 1.03E-02; chr8:23023210: effect size = 0.912, FDR = 1.37E-02, chr8:23023192: effect size = 0.926, FDR = 1.73E-02) and bisulfite pyrosequencing (chr8:23023192: t-test = 8.28E-3; chr8:23023240: t-test = 1.3E-2) of the DMPs annotated to *Ank1*. Black and blue dots refer to WT and rTg4510 TG respectively.

Strikingly, eight genes (*Nrxn2, St7, Prdm16, Camta1, Lhpp, Runx2, Map7,* and *Runx3*) were commonly associated with DMPs in all three datasets: tau pathology in rTg4510 entorhinal cortex, Aβ pathology in J20 entorhinal cortex, and human AD cortex (**Table S8, Table S9**). Many of these DMPs were annotated to the same homologous intronic region of *Prdm16/PRDM16* (rTg4510 entorhinal cortex: 10 DMPs; J20 entorhinal cortex: 2 DMPs) corresponding to a region of differential DNA methylation associated with human AD (hg19: chr1:2995301 - chr1:3332264) (**Figure S9**). Importantly, *PRDM16* is located adjacent to *PRDM16-DT*, a long non-coding RNA critical for maintaining astrocyte homeostasis and known to be downregulated in AD cortex^46^.

Together, these findings demonstrate a significant convergence of DNAm alterations between transgenic mouse models and human AD brain, reinforcing the utility of these models for studying disease-relevant epigenetic mechanisms and highlighting shared regulatory loci - including *ANK1* and *PRDM16* - that may underpin pathophysiological processes across species.

## Discussion

In this study, we identified DNAm changes in the entorhinal cortex and hippocampus associated with the progression of AD-associated neuropathology in TG models of both tau (rTg4510) and Aβ (J20). To our knowledge, this study represents the most systematic analysis of brain epigenetic variation in models of both tau and Aβ pathology. By profiling a large number of samples spanning multiple time points that encompass the development of AD pathology, integrating RRBS and methylation array technologies, and comparing changes across brain regions and with human epigenomic data, we uncover robust epigenetic signatures of neurodegeneration that offer insight into the early molecular changes associated with pathology.

Our findings suggest that tau and Aβ progression are associated with widespread and partially distinct DNAm alterations, consistent with our prior transcriptomic work in the same models. In the rTg4510 tauopathy model, we observed a substantial number of DMPs, including both expected changes linked to the *MAPT* transgene and novel alterations in genes implicated in neuronal function (e.g. *Creb314*, *As3mt, Zfp423*), ubiquitin signaling (e.g. *Dcaf5*), and apoptosis regulation. Functional enrichment analysis linked these epigenetic changes to biological processes known to contribute to tau-mediated neurotoxicity, including apoptosis and lipid signaling pathways, supporting a role for DNAm in modulating disease-relevant gene networks.

In contrast, the J20 Aβ model exhibited more modest DNAm changes, with effect sizes notably smaller than those observed in rTg4510 mice. Our results reinforce the notion that Aβ pathology engages distinct molecular pathways compared to tau pathology. Nonetheless, we identified several J20 DMPs annotated to genes with established roles in AD (e.g. *Grk2*, *Ncam2*, *Zmiz1,* and *Prmt8*). Notably, many of these genes are involved in synaptic organization and mitochondrial homeostasis, and the strong enrichment of J20 DMPs in immune-related pathways aligns with prior transcriptomic studies^47^ and further underscores the importance of neuroinflammation in Aβ-driven disease progression.

Our results suggest that the extent of DNAm alterations associated with pathology is larger in the hippocampus compared to the entorhinal cortex, particularly in the rTg4510 model. While the entorhinal cortex is among the earliest regions affected in AD, our data suggest that hippocampal pathology may drive more pronounced epigenetic dysregulation, potentially reflecting cumulative effects of pathology over time and neurodegeneration. The observation that tau-associated DNAm changes were more regionally consistent than Aβ-associated changes suggests that tau may exert a broader, more uniform influence on the epigenome, while Aβ-related effects may be more spatially constrained, slower to develop or context-dependent.

Importantly, a subset of the DNAm differences we identified in mouse models mirrored those reported in human AD postmortem cortex. This cross-species concordance strengthens the translational relevance of our findings. For example, hypermethylation of *Ank1*, previously identified as a robust epigenetic signature of tau pathology in human AD^45^, was recapitulated in rTg4510 mice and validated through bisulfite pyrosequencing. Similarly, consistent hypermethylation of the *Prdm16* locus across mouse models and human brain samples highlights conserved epigenetic responses to neuropathology and supports the utility of these models for dissecting pathogenic mechanisms.

Despite the strengths of this study, some limitations must be acknowledged. First, the RRBS and array-based approaches used in this study primarily target CpG rich regions of the genome, and do not provide information about DNAm at a large proportion of sites in the genome.

Second, while DNAm changes associated with the transgenes provide mechanistic insights, some likely reflect direct effects of transgene insertion or unrelated developmental perturbations. We attempted to mitigate this by focusing on pathology-associated DMPs and validating results using multiple platforms. Third, although mouse models recapitulate key features of AD, they lack the full complexity of human disease. Fourth, our analysis was performed on ‘bulk’ entorhinal cortex and hippocampal tissue, comprising a mix of different neural cell types. Given evidence of cell-specific epigenetic signatures in AD and changes in the proportion of different cell-types associated with the development of pathology, this may influence our ability to robustly detect pathology-associated variation. Fifth, our analyses were restricted to CpG methylation, while future studies incorporating other epigenetic layers (e.g. hydroxymethylation, chromatin accessibility) will offer a more complete picture of regulatory changes in AD. Finally, to minimize heterogeneity in our study, we further profiled only female mice, though a number of sex differences have been previously reported for these models, with females demonstrating elevated and more progressive pathology than males^48^. Future work would cross-examine results from our study with epigenetic variation in other mouse models, tissue types and male mice for a more comprehensive understanding of the development of tau and amyloid pathology in the AD brain.

In conclusion, our study highlights widespread changes in DNAm associated with the accumulation of both tau and Aβ, and supports the hypothesis that epigenetic alterations play a key role in progression of disease processes. By identifying both model-specific and conserved DNAm signatures associated with tau and Aβ pathology, we highlight genes and pathways (such as *SATB1* involved in inhibiting cellular senescence and *PRDM16* important in regulating astrocyte function) that may provide mechanistic insight and represent potential therapeutic targets. Future work integrating longitudinal epigenetic, transcriptomic, and proteomic data will be critical for elucidating the causal role of these changes in AD pathogenesis and progression.

## Methods

### Experimental model and samples

Entorhinal cortex and hippocampal tissue was dissected from rTg4510 and J20 transgenic (TG) mice and age-matched wild-type littermates (WT), in accordance with the UK Animals (Scientific Procedures) Act 1986 and approved by the local Animal Welfare and Ethical Review Board.

Details of breeding, housing conditions, and sample preparation, including neuropathology quantification, are provided in Castanho et al. (2020)^16^. Entorhinal cortex and hippocampal tissue was dissected from 31 female WT mice and 30 female rTg4510 TG, aged 2, 4, 6 and 8 months (n = 7 - 8 mice per group); and 32 female WT mice and 31 female J20 TG aged 6, 8, 10 and 12 months of age (n = 9 - 10 mice per group). Two hippocampus samples failed QC due to low signal intensity, and were subsequently removed from analysis. Only female mice were used in this study because they exhibit more pronounced pathology compared to males^48,49^, and to minimize heterogeneity between samples.

### DNA isolation

Samples were anonymized, randomized, and processed in blinded batches. Genomic DNA was extracted from the entorhinal cortex and hippocampus using the AllPrep DNA/RNA Mini Kit (Qiagen) and quantified for purity with a NanoDrop 8000 spectrophotometer (Thermo Scientific).

### Reduced Representation Bisulfite Sequencing (RRBS)

RRBS libraries were prepared using the Premium RRBS kit (Diagenode) with input concentrations measured using the Qubit dsDNA HS Assay (Thermo Fisher). Sample pooling was based on qPCR quantification (StepOnePlus Real-Time PCR System, Thermo Fisher), and bisulfite-converted pools were quantified with the NEBNext Library Quant Kit (New England Biolabs) to determine enrichment PCR cycle number. Library quality was assessed with High Sensitivity D1000 ScreenTape on a 2200 TapeStation (Agilent), then sequenced on an Illumina HiSeq2500 using 50 bp single-end reads as previously described^50^. Raw FASTQ files, post quality control (QC) assessment using FASTQC with all samples passing QC, were trimmed with *Trim Galore* (v0.4.5, Phred score = 20), aligned to the mouse reference genome (GRCm38.p4, mm10) with *Bismark* (v0.19.0) (--merge_non_CpG --comprehensive) and *Bowtie2* (v2.3.4.1) to generate coverage files. Methylation coverage across adjacent palindromic CpG sites were merged using *coverage2cytosine* (--merge_CpG --discordance 100). Bismark output values across all the samples were subsequently processed using *BiSeq* (v1.26.0). Raw methylation data was filtered by coverage (minCov = 10, perc.samples = 0.5) with remaining CpG sites smoothed to reduce weighting from sites with unusually high coverage.

### Mammalian methylation array

Microarray-based DNA methylation data (entorhinal cortex and hippocampus) was generated using a custom Illumina methylation array (HorvathMammalMethylChip40) covering 37,492 sites^14^. DNA bisulfite conversions were performed using the Zymo EZ DNA Methylation Kit (ZymoResearch, USA). DNAm levels (β values) were determined by calculating the ratio of intensities between methylated (signal A) and unmethylated (signal B) sites, ranging from 0 (completely unmethylated) to 1 (completely methylated). QC assessment was performed to remove samples with low signal intensities, low bisulfite conversion rate (80%) and a mismatch between predicted (determined by sex probe intensities) and reported sex. The *SeSaMe*^51^ method was subsequently used to normalize β values of the remaining samples for each probe.

### Differential DNA methylation analysis

Differentially methylated positions (DMPs) in the RRBS and array datasets were identified using a beta regression analysis performed with the R package *betareg*. As described in our previous study^16^, significant genotype effects between WT and TG mice and interaction effects associated with progressive changes across age between WT and TG mice were identified by testing the full model (∼ genotype + age + genotype*age) against the null model (∼ genotype + age) for the RRBS dataset, and the likelihood ratio test were used with the model (∼ genotype + age + genotype*age + Chip ID) for the array dataset. To detect significant pathology effects indicating progressive changes in TG mice, the likelihood ratio test (LRT) was used with the model: ∼ pathology (RRBS) or ∼ pathology + chip ID (array) with the quantification of tau or Aβ from immunohistochemistry staining as a measure of pathology^16^. Effect sizes and direction were deduced from the beta estimate of the regression analysis. Gene-wide P values were generated by combining probe-level P values using the Empirical Brown’s method^52^. The Pearson’s r correlation coefficient is reported unless stated otherwise.

Significant tissue effects between the entorhinal cortex and hippocampus in WT and TG mice from the array datasets were identified using the model: ∼ tissue + genotype + tissue*genotype + age + (1|Mouse). Differentially methylated positions were subsequently annotated using *ChIPseeker* (v1.22.1, parameters: annoDb = "org.Mm.eg.db", tssRegion = c(–1500, 1500)). Probes annotated to the exonic and intron regions of the gene were further cross-referenced and annotated using the mouse reference annotation (vM22, GENCODE) or the mammalian methylation manifest file. In identifying AD-associated epigenetic signatures in the entorhinal cortex where we profiled DNA methylation using RRBS and the mammalian array, the significant DMPs from the two separate differential methylation analyses were combined and treated as one dataset.

### Gene ontology

Genomic Regions Enrichment of Annotations Tool (GREAT) was used to perform functional enrichment analysis of the differentially methylated CpG sites associated with tau and Aβ pathology in the rTg4510 and J20 entorhinal cortex, respectively. We imported the bed file of the corresponding mouse coordinates (mm10) to the GREAT web service (v4.04). To account for reduced genome representation, we used a background bed file that included: (i) CpG sites detected using RRBS and annotated to the promoter using *ChIPseeker*, (ii) all differentially expressed CpG sites identified in the rTg4510 and J20 TG vs WT cortex and across pathology, and (iii) all the CpG sites in the mammalian methylation array.

### Comparative analysis with human data

Differentially methylated positions associated with tau pathology in a meta-analysis of human AD data were extracted from Supplementary Data 8 from our recent publication^15^. For cross-species comparisons, the human UCSC gene annotations were subsequently converted to the equivalent homologous mouse gene names, considering only mouse-specific homologous genes. This gene set was then used to determine the overlap of genes annotated with DMPs identified in the current study.

### Validation of DNA methylation changes using bisulfite pyrosequencing

Forward and reverse primers targeting CpG sites annotated to *Ank1* (chr8: 23023193 - 23023241, mm10) were designed using PyroMark Assay Design software 2.0 (Qiagen, UK). DNA from rTg4510 mice (n = 14, 2 and 8 months) was isolated using AllPrep DNA/RNA Mini Kit (QIAGEN) and underwent bisulfite conversion using the EZ DNA Methylation-Gold™ Kit (Zymo Research). Following PCR amplification, DNA methylation was profiled using the PyroMark Q48 AutoPrep platform (Qiagen, UK).

### Data and code availability

Raw mammalian methylation array data and RRBS FASTQ for both rTg4510 and J20 have been deposited in GEO under accession number GSE246561. Intermediate files are available on Zenodo (doi: 10.5281/zenodo.15741353). All code supporting this study is available at https://github.com/SziKayLeung/AD_mouse_methylation.

## Acknowledgments

This work was funded in part through the Medical Research Council (MRC) Proximity to Discovery: Industry Engagement Fund (Precision Medicine Exeter Innovation Platform reference MC_PC_14127) and a research grant from Alzheimer’s Research UK (ARUK-PG2018B-016).

I.C.’s doctoral studentship was supported by the Alzheimer’s Society in partnership with the Garfield Weston Foundation (grant 231). Sequencing and computational facilities were supported by an MRC Clinical Infrastructure award (MR/M008924/1); the Wellcome Trust Institutional Strategic Support Fund (WT097835MF); a Wellcome Trust Multi-User Equipment Award (WT101650MA); and a Biotechnology and Biological Sciences Research Council (BBSRC) Longer and Larger (LoLa) award (BB/K003240/1).

## Author Contributions

J.M., Z.A., and I.C designed the study. I.C conducted RRBS, array, and immunohistochemistry laboratory experiments. S.K.L. led primary data analysis and bioinformatics, with analytical and computational input from I.C. (RRBS pre-processing), E.W. (RRBS differential methylation analysis), A.D. (array pre-processing and analysis), D.S.V. (DMP annotations), and E.H. J.M., K.L., and E.D. obtained funding. S.P. and R.S. performed pyrosequencing under the supervision of A.R.S. S.K.L, J.M., and I.C. drafted the manuscript. All authors read and approved the final submission.

