## Supplementary Tables Legend for "Methylomic signatures of tau and amyloid-beta in transgenic mouse models of Alzheimer’s disease neuropathology"

**Supplementary Table 1. Phenotypic information for the mouse samples sequenced in this study.**
Neuropathology refers to Castanho et al. 2020; J20 amyloid, rTg4510 tau. “X” in ECX and HIP array column refers to the samples that were sequenced using the Illumina mammalian DNA methylation array. The burden of tau or amyloid neuropathology is expressed as percentage area.

**Supplementary Table 2. Differentially methylated sites associated with tau pathology in rTg4510 TG and WT mice, and associated with progressive tau pathology.**
Tabulated are the sites identified as significantly differentially methylated (FDR < 0.05) between rTg4510 TG and WT mice (i.e. genotype effect) and in rTg4510 TG mice associated with tau pathology (i.e. pathology effect) using RRBS and array profiling. Sites are further classified as TRUE if identified as significant with an interaction effect. Corresponding CpG probes are provided if sites were detected using illumina mammalian methylation array.

**Supplementary Table 3. Gene ontology analysis**
Tabulated are the results from functional enrichment analysis of the differentially methylated CpG sites associated with tau and amyloid pathology in the rTg4510 and J20 entorhinal cortex respectively.

**Supplementary Table 4. Differentially methylated sites associated with amyloid-beta pathology in J20 TG and WT mice, and associated with progressive amyloid-beta pathology.**
Tabulated are the sites identified as significantly differentially methylated (FDR < 0.05) between J20 WT and TG mice (i.e. genotype effect), and in J20 TG mice associated with amyloid-beta pathology, using RRBS and array profiling. Corresponding CpG probes are provided if sites were detected using illumina mammalian methylation array.

**Supplementary Table 5. Common DMPs associated with tau and beta-amyloid pathology in rTg4510 and J20 ECX.**

**Supplementary Table 6. Differentially methylated sites associated with tau pathology in the rTg4510 hippocampus**
Tabulated are the sites identified as significantly differentially methylated (FDR < 0.05) between rTg4510 TG and WT mice (i.e. genotype effect), and with progressive tau pathology in rTg4510 TG mice in the hippocampus from array profiling. ECX column “TRUE” refers to sites also identified as significantly differentially methylated in the entorhinal cortex from array profiling (across all statistical models testing for genotype and progressive pathology effects). Genotype effect column “TRUE” and Interaction effect column “TRUE” refer to sites also significantly altered from statistical models testing for genotype and interaction effect in the rTg4510 hippocampus.

**Supplementary Table 7. Differentially methylated sites associated with Aβ pathology in the J20 hippocampus**
Tabulated are the sites identified as significantly differentially methylated (FDR < 0.05) between J20 TG and WT mice (i.e. genotype effect), and with progressive Aβ pathology in J20 TG mice in the hippocampus from array profiling. ECX column “TRUE” refers to sites also identified as significantly differentially methylated in the entorhinal cortex from array profiling (across all

statistical models testing for genotype and progressive pathology effects). Genotype effect column "TRUE" and Interaction effect column "TRUE" refer to sites also significantly altered from statistical models testing for genotype and interaction effect in the J20 hippocampus.

**Supplementary Table 8. Overlap of genes with DMPs associated with tau pathology in rTg4510 cortex and AD-associated pathology in human *post-mortem* cortex**
Tabulated are common genes annotated with significantly differentially methylated (FDR < 0.05) sites between rTg4510 TG and WT mice (i.e. genotype effect), and with progressive tau pathology in rTg4510 TG mice in the cortex from RRBS and array profiling. HumanTauPathology column "TRUE" refers to genes that were identified with sites as significantly differentially methylated in the human *post-mortem* cortex associated with tau (measured using Braak staging) pathology. The human DMPs associated with tau pathology are taken from the cross-cortex meta-analysis at Bonferroni significance (P < 1.24E-07)[1].

**Supplementary Table 9. Overlap of genes with DMPs associated with Aβ pathology in J20 cortex and AD-associated pathology in human *post-mortem* cortex**
Tabulated are common genes annotated with significantly differentially methylated (FDR < 0.05) sites between J20 TG and WT mice (i.e. genotype effect), and with progressive Aβ pathology in J20 TG mice in the cortex from RRBS and array profiling. HumanAmyloidPatthology column "TRUE" refers to genes that were identified with sites as significantly differentially methylated in the human *post-mortem* cortex associated with amyloid-beta pathology (measured using Thal phase). The human DMPs associated with tau pathology are taken from the cross-cortex meta-analysis at Bonferroni significance (P < 1.24E-07)[1].
