## Supplementary Figures for "Methylomic signatures of tau and amyloid-beta in transgenic mouse models of Alzheimer’s disease neuropathology"


| | |
|---|---|
| Supplementary Figure 1 | Annotations of DNA methylation sites from RRBS profiling |
| Supplementary Figure 2 | Methylation of common sites profiled using RRBS and array |
| Supplementary Figure 3 | DMPs annotated to *Mapt, Prn/Prnp, Ncapg2, Fgf14* |
| Supplementary Figure 4 | Pyrosequencing validation of *Prn/Prnp* DMPs |
| Supplementary Figure 5 | Correlation of rTg4510 ECX vs HIP effect sizes |
| Supplementary Figure 6 | Common DMPs associated with tau pathology in ECX & HIP |
| Supplementary Figure 7 | Correlation of J20 ECX vs HIP effect sizes |
| Supplementary Figure 8 | Epigenetic age determined using calibrated epigenetic clock |
| Supplementary Figure 9 | Differential methylation changes annotated to Prdm16/PRDM16 |

**Supplementary Figure 1: Annotations of DNA methylation sites from RRBS profiling**
Summary of ChIPseeker annotations of all DNA methylation sites profiled using RRBS in rTg4510 (n = 31 WT, n = 30 TG) and J20 entorhinal cortex tissue (n = 32 WT, n = 31 TG).

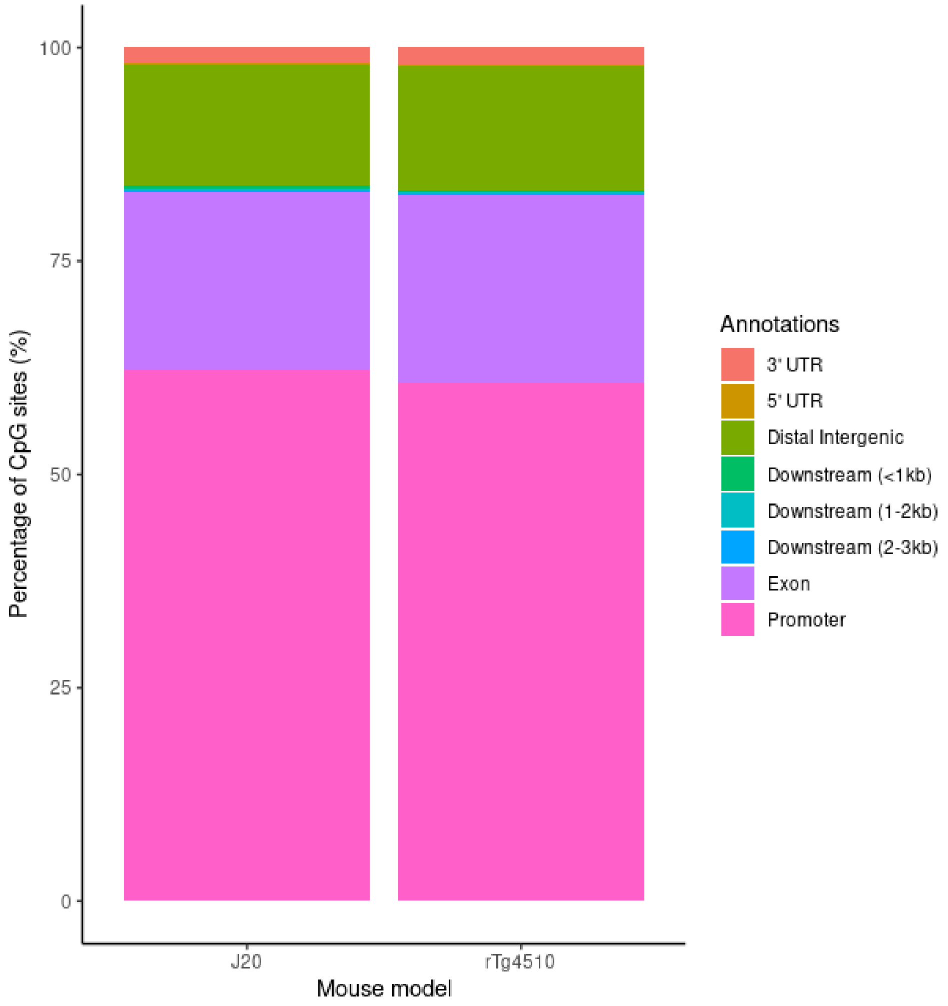

**Supplementary Figure 2: There is a high correlation of DNA methylation levels measured using RRBS and Illumina DNAm arrays for sites profiled using both technologies.** Shown are density plots of the mean DNA methylation values (determined after *BiSeq* processing) of all overlapping CpG sites profiled using the mammalian methylation array and RRBS in the **(A)** rTg4510 entorhinal cortex (n = 61), and **(B)** J20 entorhinal cortex (n = 63).

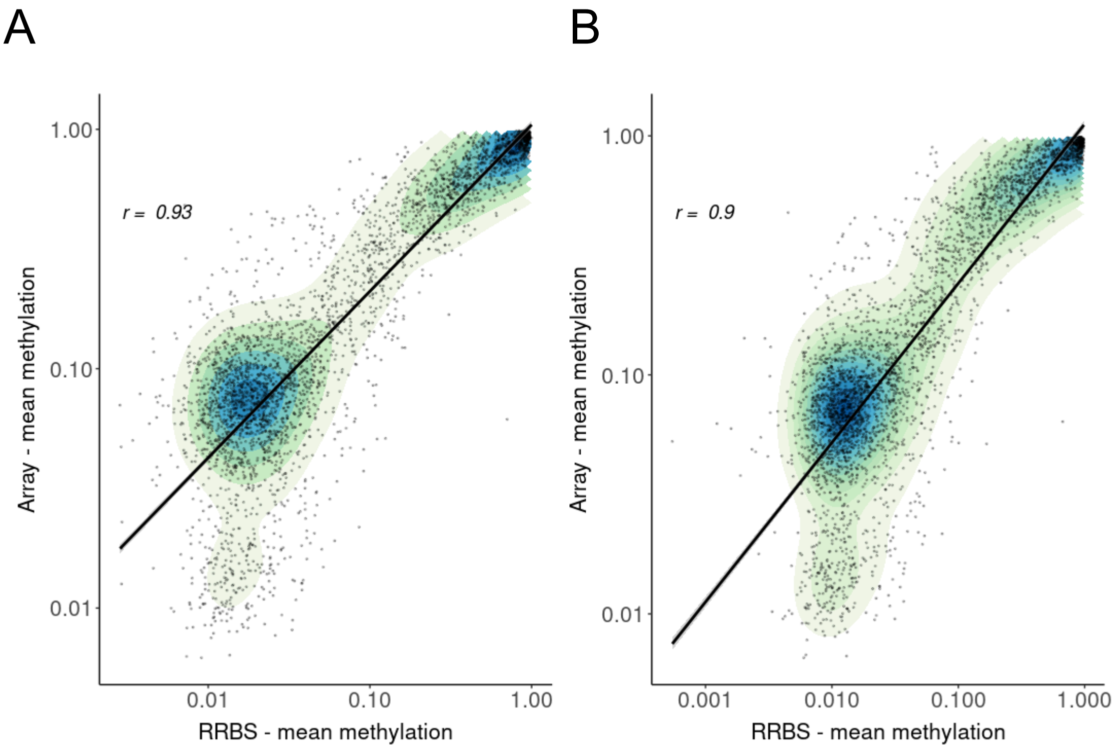

**Supplementary Figure 3: Differentially methylated positions annotated to *Mapt, Prn/Prnp, Ncapg2, Fgf14* as a consequence of transgene insertion in rTg4510 TG mice.**
Shown are the gene tracks and differentially methylated sites annotated to **(A)** *Mapt* (chr11:104318231, effect size = -1.04, FDR = 3.73E-30), **(B)** *Prn/Prnp* (n = 4 CpG sites, chr2:131910162 - chr2:131910201, mean effect size = 1.15), **(C)** *Ncapg2* (chr12:116425797, effect size = -0.19, FDR = 1.17E-2) and **(D)** *Fgf14* (chr14:124676565, effect size = 0.54, FDR = 1.89E-6), between WT and rTg4510 TG mice. Black and blue dots refer to WT and TG respectively.

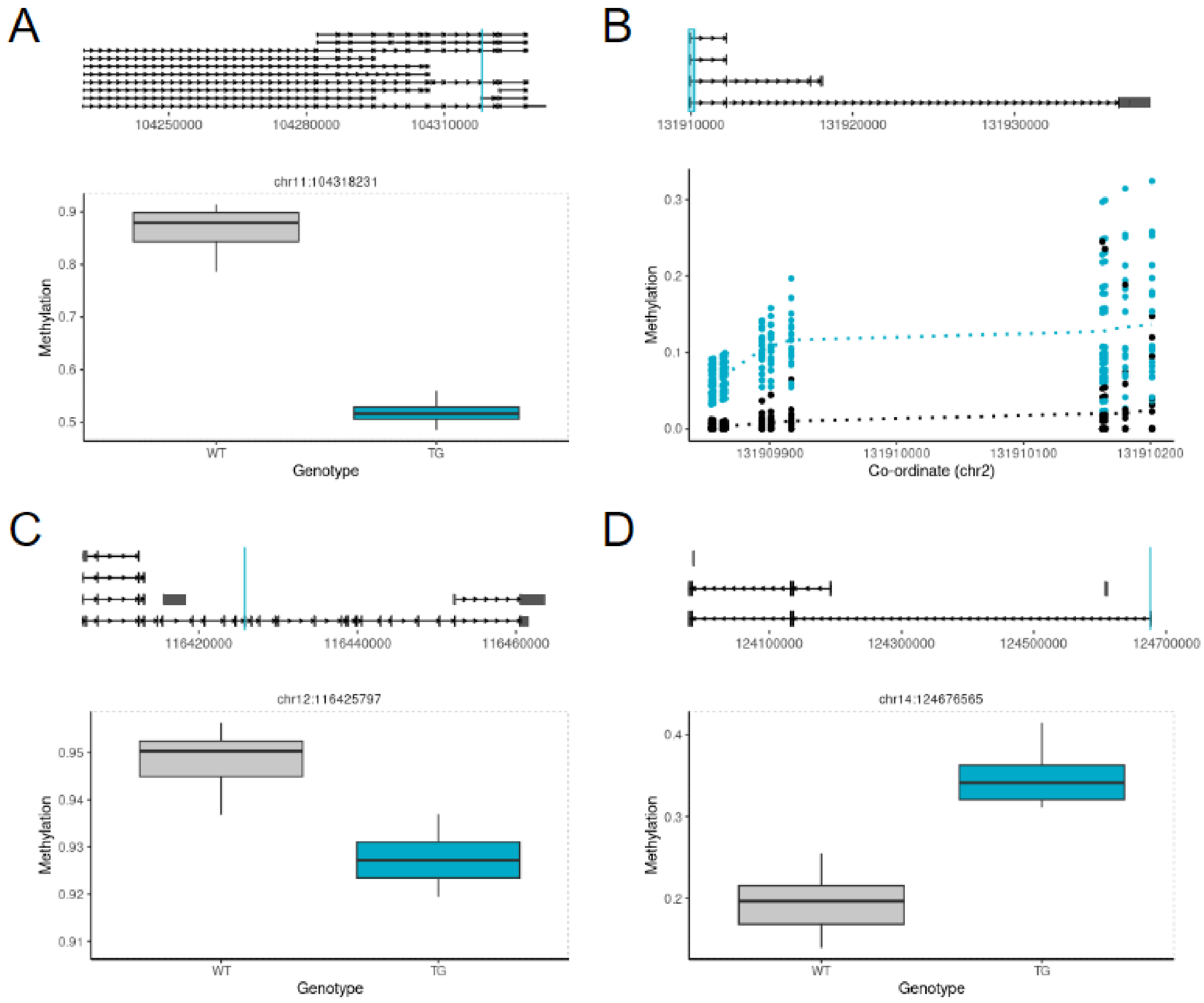

**Supplementary Figure 4: Pyrosequencing validation of *Prn/Prnp* DMPs.**
Shown is a scatter plot of the DNA methylation estimates determined from bisulfite pyrosequencing of a subset of DMPs annotated to *Prn/Prnp* in the rTg4510 entorhinal cortex (chr2:131910162: t-test P = 4.31E-25; chr2:131910164: t-test P = 6.70E-33; chr2:131910180: t-test P = 1.80E-36; chr2:131910201: t-test P = 1.59E-39), validating the results observed from RRBS (n = 4 CpG sites, chr2:131910162 - chr2:131910201, mean effect size = 1.15, Empirical Brown's method: P = 7.00E-5).

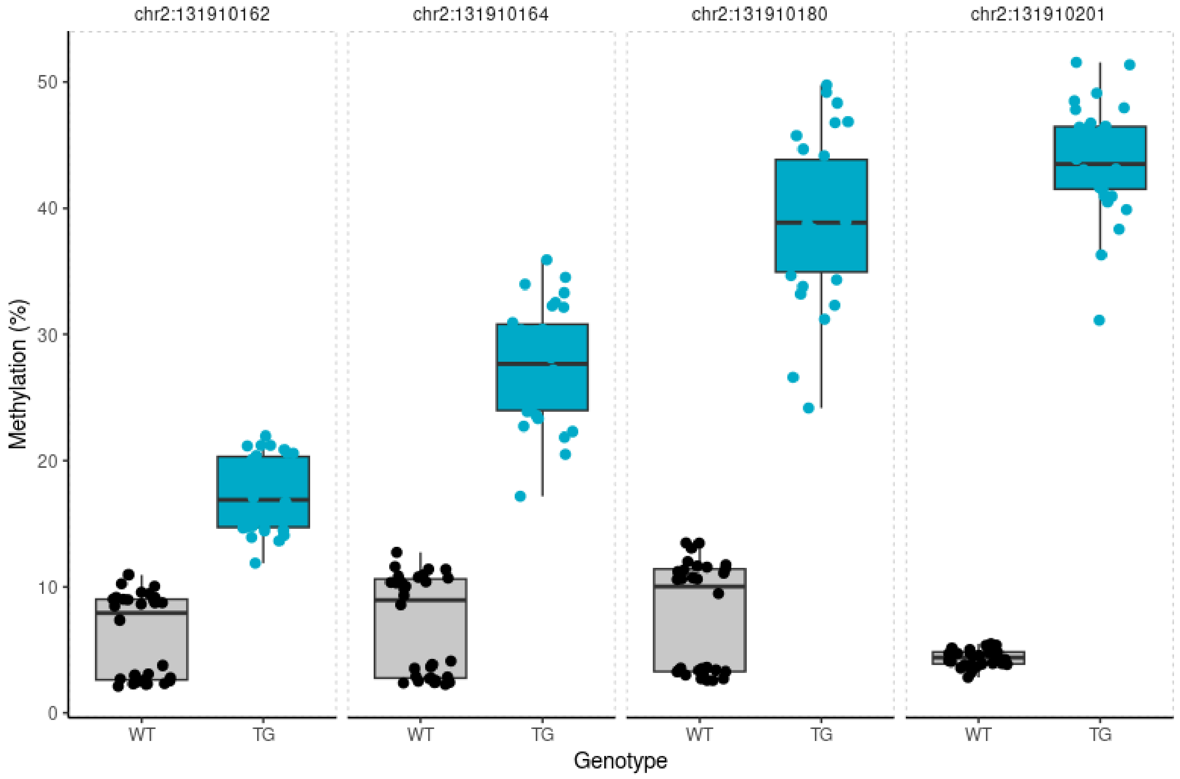

**Supplementary Figure 5: Pathology- but not genotype-associated DNAm differences are strongly correlated between entorhinal cortex and hippocampus in rTg4510 mice.**
For all sites profiled in both the entorhinal cortex and hippocampus using the Illumina DNA methylation array, we observed that **(A)** genotype-associated differences are not correlated across brain areas but **(B)** pathology-associated differences are highly correlated across tissues. **(C)** and **(D)** show the same relationships for significant DMPs identified in both brain areas. Pearson's correlations are provided. ECX – entorhinal cortex, HIP – hippocampus.

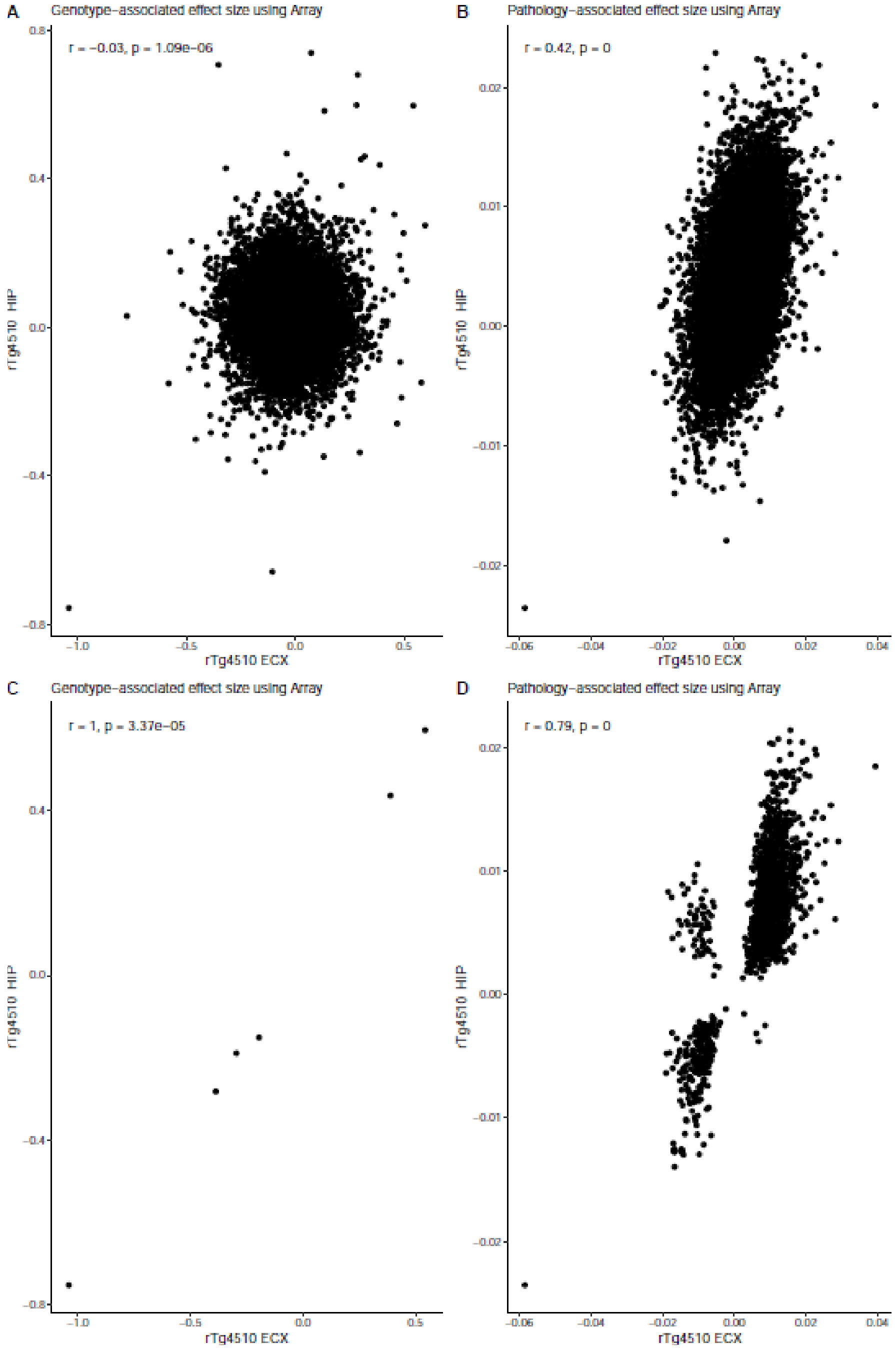

**Supplementary Figure 6: Common differentially methylated sites between TG and WT rTg4510 mice between the entorhinal cortex and hippocampus.**

Shown are DMPss annotated to **(A)** *Dcaf5* (chr12:80436248), **(B)** *Satb1* (chr17:51746925), **(C)** *Cltc* (chr11:8670046), **(D)** *Mapt* (chr11:104318231), **(E)** *Ncapg2* (chr12:116425797), and **(F)** *Fgf14* (chr14:124676565) associated with genotype in rTg4510 TG mice in the entorhinal cortex (ECX) and hippocampus (HIP). Black and blue dots refer to WT and TG, respectively.

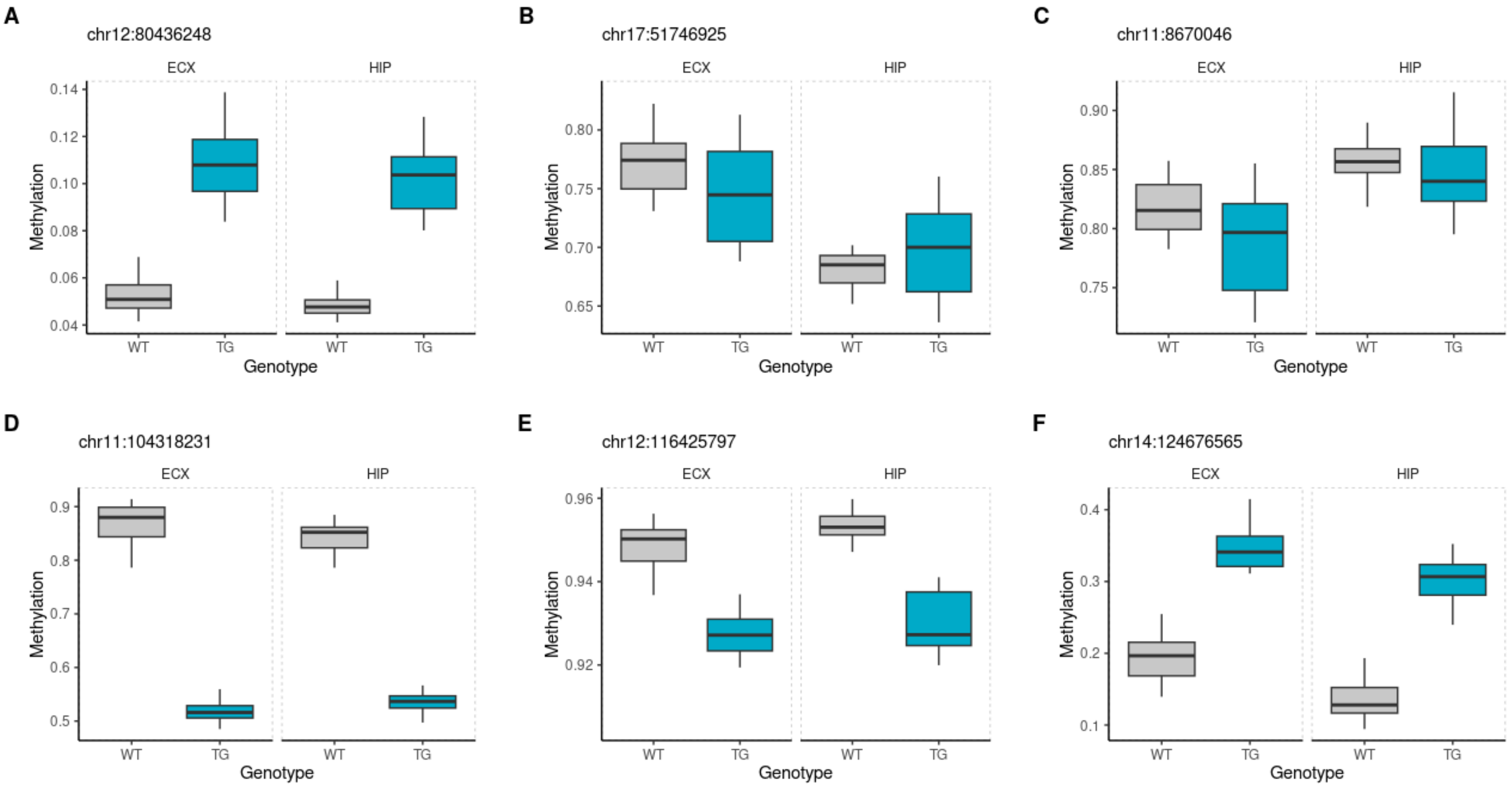

**Supplementary Figure 7: No correlation of genotype- and pathology-associated DNAm differences between entorhinal cortex and hippocampus in J20 mice.**
For all sites profiled in both the entorhinal cortex and hippocampus using the Illumina DNA methylation array, we observed no correlation of **(A)** genotype-associated differences, and **(B)** pathology-associated differences across brain areas across tissues. No common significant sites associated with pathology were identified between J20 cortex and hippocampus. ECX – entorhinal cortex, HIP – hippocampus.

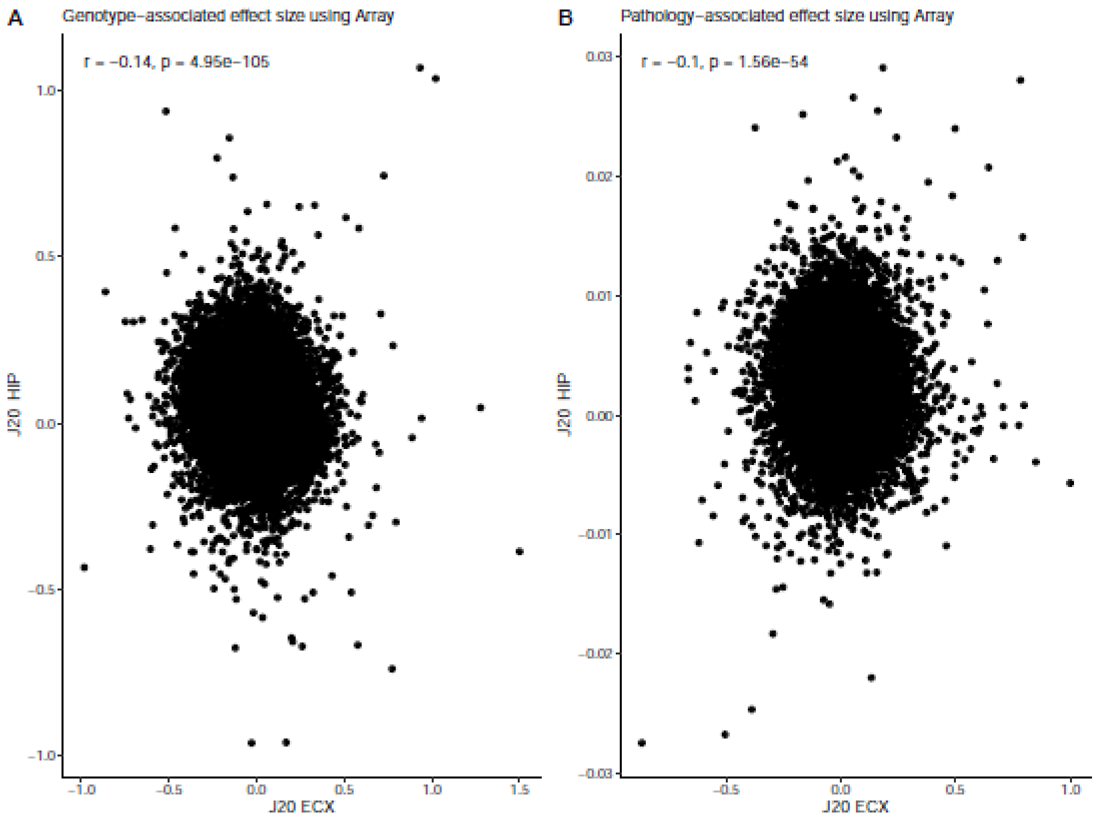

**Supplementary Figure 8: Epigenetic age determined using epigenetic clock calibrated on mouse cortex.**

Shown are scatter plots of the epigenetic age of the **(A)** rTg4510 entorhinal cortex (r = 0.89, P = 1.52E-14), **(B)** rTg4510 hippocampus (r = 0.87, P = 7.39E-14), and **(C)** J20 entorhinal cortex (r = 0.81, P = 2.1E-10) and **(D)** J20 hippocampus (J20: r = 0.90, P = 4.65E-15). The epigenetic age is derived from DNA methylation determined using the mammalian methylation array previously calibrated using the mouse cortex.

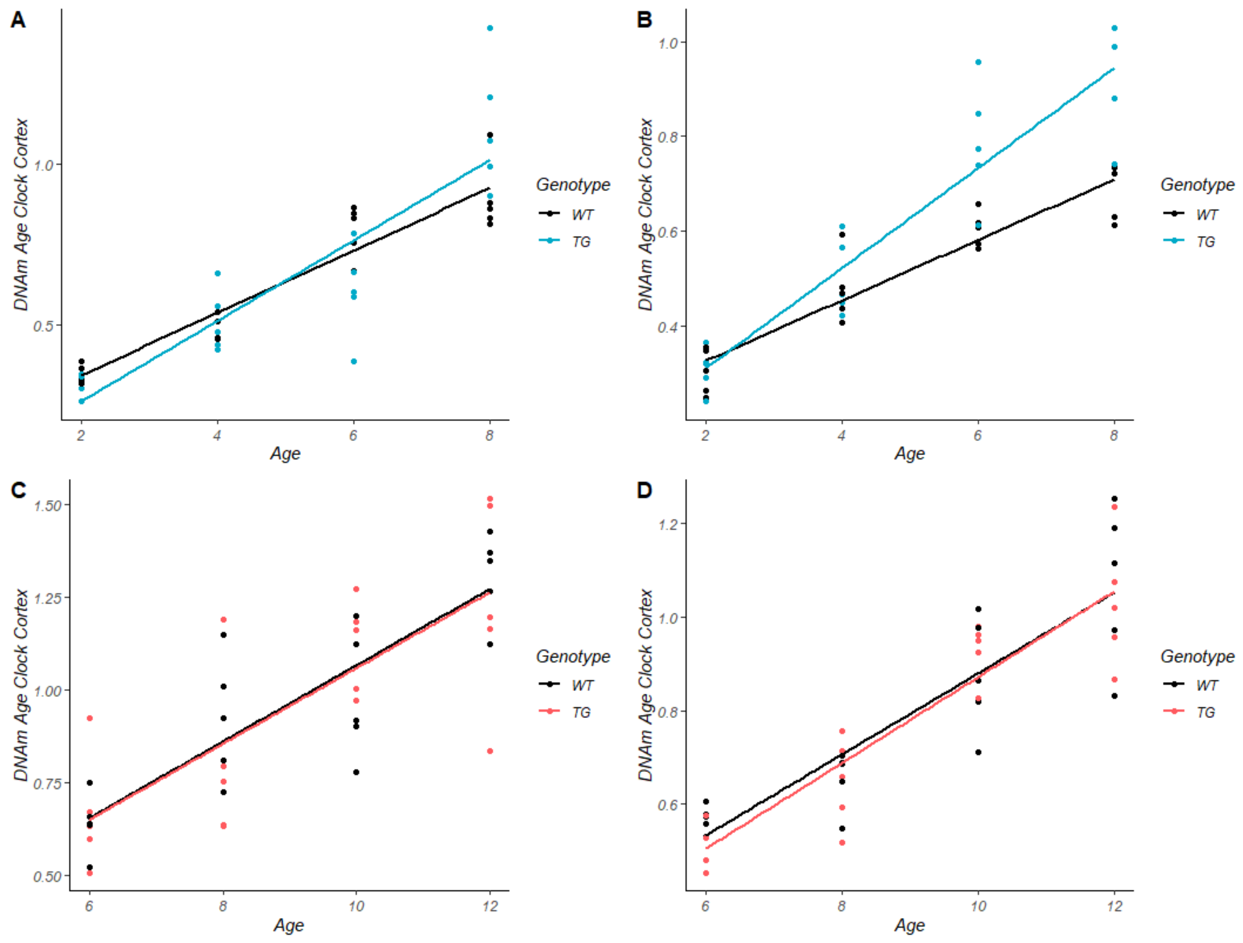

**Supplementary Figure 9: Differential methylation changes annotated to *Prdm16/PRDM16* in rTg4510 ECX, J20 ECX and AD *post-mortem* cortex.**

Shown are the gene tracks and differentially-methylated positions annotated to *Prdm16* **(i)** between WT and rTg4510 TG mice (chr4:154640585, effect size = 1.42, FDR = 3.49E-2; chr4:154640557, effect size = 1.41, FDR = 3.97E-2; chr4:154346846, effect size = -1.42, FDR = 4.97E-2), and **(ii)** WT and J20 TG mice (chr4:154519364, effect size = 2.12, FDR = 3.86E-2). Black, blue and red dots refer to WT, rTg4510 TG, J20 TG respectively, and lines on the tracks refer to the location of CpG sites.

i

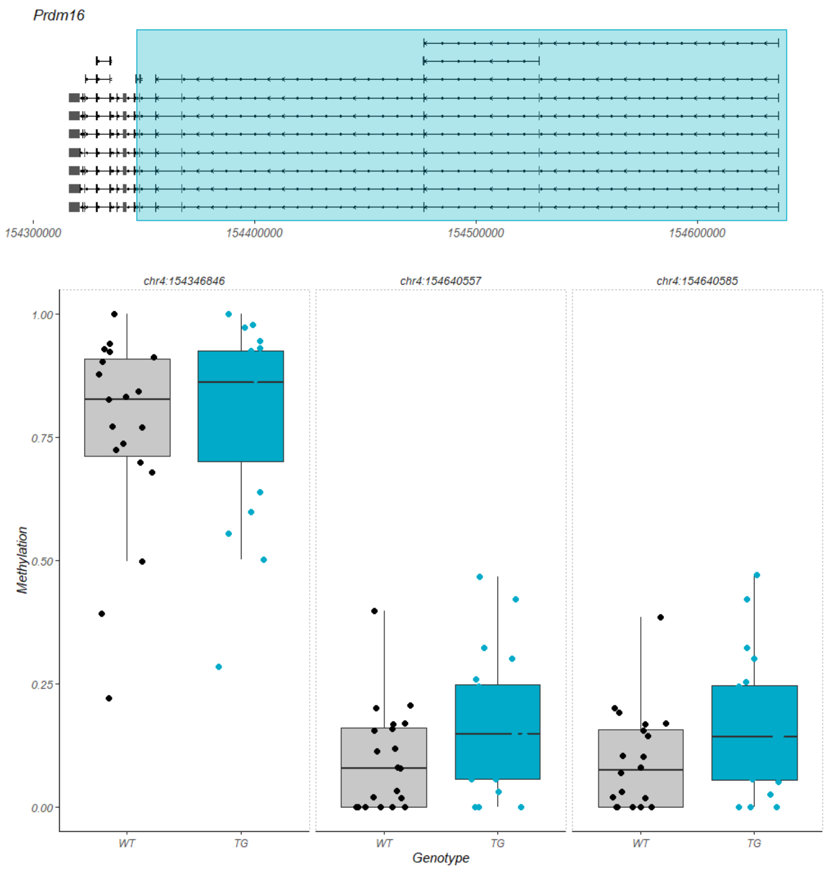


ii

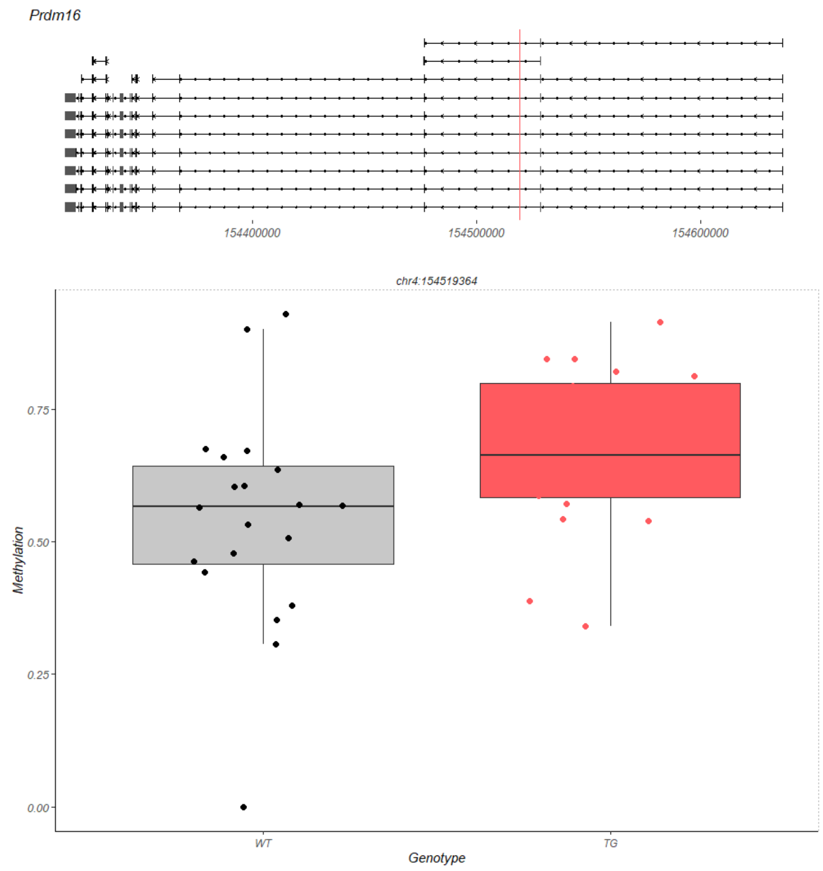
